# The choline-binding proteins PspA, PspC and LytA of *Streptococcus pneumoniae* and their role on host cellular adhesion and damage

**DOI:** 10.1101/2022.07.08.499412

**Authors:** Cláudia Vilhena, Shanshan Du, Miriana Battista, Martin Westermann, Thomas Kohler, Sven Hammerschmidt, Peter F. Zipfel

**Affiliations:** Department of Infection Biology, Leibniz Institute for Natural Product Research and Infection Biology, Jena, Germany; Centre for Electron Microscopy, Jena University Hospital, Friedrich-Schiller-University of Jena, Jena, Germany; Department of Molecular Genetics and Infection Biology, Interfaculty Institute for Genetics and Functional Genomics, Center for Functional Genomics of Microbes, University of Greifswald, Greifswald, Germany; Institute of Microbiology, Friedrich-Schiller-University, Jena, Germany

**Author notes:** **Corresponding author**: Prof. Dr. Peter F. Zipfel, Department of Infection Biology, Leibniz Institute for Natural Product Research and Infection Biology, Hans Knöll Institute, Beutenbergstrasse 11a, 07745 Jena, Germany.

**Keywords:** biofilms, hemolysis, endothelial, cell wall, immune evasion, hemolytic uremic syndrome

## Abstract

*Streptococcus pneumoniae* is a Gram-positive opportunistic pathogen that can colonize the upper respiratory tract. It is a leading cause of a wide range of infectious diseases, including community-acquired pneumonia, meningitis, otitis media and bacteraemia. Pneumococcal infections cause 1–2 million deaths per year, most of which occur in developing countries, where this bacterial species is probably the most important pathogen during early infancy. Here, we focused on choline-binding proteins (CBPs), i.e., PspC, PspA and LytA, and their integration into and interaction with the cell wall of *S. pneumoniae*. The three pneumococcal proteins have different surface-exposed regions but share related choline-binding anchors. These surface-exposed pneumococcal proteins are in direct contact with host cells and have diverse functions. PspC and PspA bind several host plasma proteins, whereas LytA plays a role in cell division and the lytic phase. We explored the role of the three CBPs on adhesion and pathogenicity in a human host by performing relevant imaging and functional analyses, such as electron microscopy, confocal laser scanning microscopy and functional quantitative assays targeting biofilm formation and the haemolytic capacity of *S. pneumoniae*. *In vitro* biofilm formation assays and electron microscopy experiments were used to examine the ability of knockout mutant strains lacking the *lytA*, *pspC* or *pspA* genes to adhere to surfaces. The mutant strains were compared with the *S. pneumoniae* D39 reference strain. We found that LytA plays an important role in robust synthesis of the biofilm matrix. PspA and PspC appeared crucial for the haemolytic effects of *S. pneumoniae* on human red blood cells. Furthermore, all knockout mutants caused less damage to endothelial cells than wild-type bacteria, highlighting the significance of CPBs for the overall pathogenicity of *S. pneumoniae*. Hence, in addition to their structural function within the cell wall of *S. pneumoniae*, each of these three surface-exposed CBPs controls or mediates multiple steps during bacterial pathogenesis.

## Introduction

***Streptococcus pneumoniae*** is a human pathogen that colonizes the upper respiratory tract and can cause otitis media, bronchitis, sinusitis, community-acquired pneumonia and sepsis (1, 2). The latest data from the World Health Organization show that pneumonia kills yearly more than 800,000 children under the age of 5 years (data from 2017), accounting for 15% of all child deaths at this age. Current *S. pneumoniae* vaccines target mostly the pneumococcal polysaccharide capsule, which acts as a physical barrier to the outside and protects the bacterium from recognition by the host immune system (3). Below the capsule, the pathogen has a thick peptidoglycan and teichoic acid-based cell wall, beneath which lies a phospholipid membrane (4). Upon contact with human immune cells, pneumococci shed the capsule and expose the cell wall to the outside environment (5, 6).

*S. pneumoniae* display several virulence proteins that are integrated into the cell wall; some of these extend to the outside (7). In addition to immune escape, several proteins have structural functions, whereas others mediate carbohydrate and sugar metabolism, or control the cell division machinery (8–11). One family of these surface proteins shares related anchor domains that bind non-covalently to phosphoryl choline moieties, which are common constituents of the peptidoglycan layer and are therefore called choline-binding proteins (CBPs). CBPs are key players in immune evasion, virulence (12) and host pathogenicity (13–15). Their pathogenic roles range from proteolytic activities, binding of human plasma regulators, complement proteins and immune components, and cell-mediated contact to inactivate human immunoglobulins prior to nasopharyngeal colonization (16–19).

Two CBPs, PspA and PspC (also termed CbpA), are surface-exposed immune evasion proteins, which bind human plasminogen, Factor H, secretory IgA, vitronectin, thromospondin, laminin, C3, etc. (20–26).

PspA and PspC share structural similarities: each have modular, unique and variable N-terminal regions, a proline rich domain that varies in size and almost identical C-terminal regions (27, 28). The role of both proteins in immune evasion has been attributed to their domain organization, and to sites that bind human plasma proteins. Another CBP is LytA, an intensely studied pneumococcal autolysin (also called *N*-acetylmuramoyl-l-alanine amidase) (29) that belongs to a widely distributed group of cell wall-degrading enzymes responsible for peptidoglycan cleavage; as such, LytA plays a crucial role in cell division (30).

Pneumococcal bloodstream infections start with colonization of the upper respiratory tract, crossing of the epithelial barrier and culminates with contact to human plasma, where the pathogen is immediately confronted and attacked by the complement system (31, 32). During nasopharyngeal colonization and recurrent otitis media in children, *S. pneumoniae* forms heterogeneous microbial communities embedded in a self-producing polysaccharide matrix called a biofilm (33–35). Pneumococci, similar to many other pathogenic bacteria, co-exist and colonize the host in their multicellular form rather than in their planktonic form (36–38). Biofilms are three-dimensional structures formed by agglomerates of bacteria embedded in a self-forming polysaccharide matrix (39). This complex structure protects the bacteria against any external dangers, such as host toxins and antibiotics. Members of the *Streptococcus* genus do form biofilms (40). However, formation of these multicellular three-dimensional structures by *S. pneumoniae* has not been studied widely (41, 42). Even though a lot of effort has been put into understanding pneumococcal biofilm formation and its role in pathogenesis, some questions remain unanswered.

Therefore, the aim of this study was to increase our knowledge of the role of PspA, PspC and LytA in adhesion and pathogenicity of *S. pneumoniae.* To this end, we generated bacterial mutants each lacking the gene encoding PspA, PspC or LytA, and compared them with the parental pathogenic reference strain *S. pneumoniae* D39 (wild-type, WT). By combining state of the art single-cell microscopy, electron microscopy and infection assays, we systematically examined the relevance of each CBP to bacterial adhesion, haemolysis and cytotoxicity.

## Material and Methods

### Bacterial strains, media, and growth conditions

The pathogenic strain *Streptococcus pneumoniae* D39 was used as reference strain (43). The D39-derivated mutants used comprised: Δ*pspA,* Δ*pspC* and Δ*cps*Δ*lytA.* All *S. pneumoniae* strains were grown in liquid Todd-Hewitt broth (Roth^®^) supplemented with yeast extract (THY) at 37°C with 5% CO_2_. Blood agar plates were prepared from Blood agar (VWR^®^) with addition of 5% defibrinated sheep blood (Thermo Scientific^®^). When required, 5 µg/mL erythromycin or 100 µg/mL kanamycin was used for selection. Growth was monitored by measuring the optical density at 600 nm (OD_600_).

### Construction of *S. pneumoniae* D39-derivated mutants

The *pspA*, *pspC* and the *lytA* deletion mutants were generated in the genetic background of the nonencapsulated strain D39Δ*cps* (44, 45). The construction of the *pspC* mutant was described earlier (25). For the *lytA* mutant, the gene region of *lytA* from a *S. pneumoniae* D39Δ*lytA* insertion deletion mutant was amplified from genomic DNA (kind gift of R. Brückner, Kaiserslautern) using primer LytA_KO_f (5’ GGTGTTATCCTTTGTGAACCTC 3’) and LytA_KO_r (5’GCAATCATGCTTTGATTCAAA 3’). The resulting, 1973 bp fragment contains 498 bp upstream of *lytA*, 38 bp from the beginning of the lytA gene, followed by ermR, 42 bp from the end of the *lytA* gene and 486 bp downstream of *lytA*. The amplified PCR fragment was used to transform D39Δ*cps* using routine protocols (44). Resulting colonies were selected on LB agar plates containing kanamycin and erythromycin and verified by PCR and agarose gel analysis. Th *pspA*-mutant was described earlier (46). Briefly, plasmid pQSH29 containing the full-length *pspA* gene of strain ATCC 11733 (serotype 2) amplified by PCR using the primer combination SH20 (5’GCGCGCGCGCGGATCCTTGAATAAGAAAAAAATGATTTTAACA 3’) and SH21 (5’CTCAGCTAATT AAGCTTGCTTAAACCCATTCACCATTGGC 3’) was used to insert the antibiotic gene cassette. The BamHI/HindIII digested PCR product was then ligated with the similarly digested vector pQE30 (Qiagen, Hilden, Germany), resulting in a 1.860 *pspA* DNA-insert. The knockout plasmid was constructed by digestion of the cloned *pspA* gene with SacI and blunt ligation with the PCR amplified erythromycin gene cassette *ermB*, resulting in plasmid pMSH5.1. This plasmid was used to transform *S. pneumoniae* strains and knockouts are verified by immunoblot analysis (46).

### Cell culture and cell harvesting

Human umbilical vein endothelial cells (HUVEC, CRL-1730) and adenocarcinomic human alveolar basal epithelial cells (A549, ATCC 107) were cultivated in Dulbecco’s modified Eagle’s medium, DMEM (BioWhittaker^®^) supplemented with 10% fetal calf serum (Biochrom®), 6 mmol/L l-glutamine (BioWhittaker^®^) and a mixture of penicillin/streptomycin (100U/100 µg/mL, Sigma®) at 37°C in the presence of 5% CO_2_. The full supplemented DMEM medium will be referred to as growth medium.

Adherent human cells were washed with pre-warmed Dulbecco’s phosphate-buffered saline (DPBS) (BioWhittaker^®^) and harvested by incubation for 10 minutes at 37°C with PBS containing trypsin/EDTA (Gibco®). Cell detachment was stopped by adding 10 mL of growth medium. After centrifugation, the pellet was resuspended in 1 mL growth medium and the cells were counted using the cell counter CASY (OLS^®^CASY).

### Static biofilm model

Pneumococci biofilms were grown in either THY or DMEM media to mid-logarithmic phase. Bacteria were washed and resuspended in the corresponding medium at a concentration of 1 x 10^6^ cells/mL). Bacterial suspensions were incubated on sterile, 18 mm round glass 1.5 H coverslips (Roth^®^) in the bottom of 24-well polystyrene plates (Thermo Scientific^®^). Exceptionally, bacterial suspensions were incubated on 12-well plates containing 12 mm round glass coverslips. The plates were incubated at 37°C with 5% CO_2_ for 48 h. The growth medium was changed every 6 h. Bacterial biofilms were either evaluated by Scanning Electron Microscopy (SEM) or visualized by Confocal Laser Scanning Microscopy (CLSM).

### Biofilm quantification

For quantification of biofilm formation, the Microtiter Dish Biofilm Formation Assay was performed with small changes (O’Toole, 2010). Pneumococci were grown overnight on solid agar blood plates. Bacteria were resuspended in PBS and diluted in THY or DMEM media to reach OD_600_ of 0.1. Bacteria were statically grown on 96-well plates to obtain biofilms (Thermo Scientific^®^), at 37°C with 5% CO_2_. At the indicated time points, the supernatant was transferred to another plate and OD_600_ measured as a read for planktonic growth. To each well of the original plate, 100 µL of a 1% crystal violet solution was added and the plate was incubated for 30 min at room temperature. After repeated washing with water, the plate was left to dry for 1h. Ethanol (95% (v/v)) was added to each well and left for 30 min at room temperature. The released crystal violet was finally transferred to a new 96-well plate and absorbance at 620 nm measured. Statistical analysis was performed using Prism version 9 for Windows (GraphPad Software, La Jolla, CA).

### Confocal Laser Scanning Microscopy (CLSM)

Bacterial viability within the biofilms was evaluated using the Bacterial Viability Stain kit (Biotium^®^) according to the manufacturer’s description. Bacteria were treated as described on static biofilm section. Planktonic bacteria were removed and the remaining biofilm layer was washed with DPBS and stained with a fluorescence red marker for dead cells, Ethidium Homodimer III (EthD-III) and a green peptidoglycan dye, wheat germ agglutinin (WGA) conjugated to CF®488A. After staining, biofilms were washed to remove unbound dyes and mounted using SlowFade Diamond® (Invitrogen^®^) mounting oil. Then the coverslip was sealed with nail polish and biofilms were evaluated by confocal laser scanning microscopy using a LSM 710 fitted with ZEN 2011 software (Zeiss GmbH).

### Scanning Electron Microscopy (SEM)

For SEM, biofilms were grown on 12-well plates containing 12 mm coverslips (Roth^®^), as described above for the static biofilm model. At designated time points, medium was aspirated. Then cells were fixed for 1h in 2.5% glutaraldehyde in sodium cacodylate buffer (0.1 M, pH 7.0) and washed three times with sodium cacodylate buffer for 20 min each. Samples were dehydrated in rising ethanol concentrations followed by critical point drying, using a Leica EM CPD300 Automated Critical Point Dryer (Leica) and finally coated with gold (25 nm) in a Safematic CCU-010 HV Sputter Coating System (Safematic). SEM images were acquired at different magnifications in a Zeiss-LEO 1530 Gemini field-emission scanning electron microscope (Carl Zeiss) at 6-8kV acceleration voltage and a working distance of 5-7 mm using an InLense secondary electron detector for secondary electron imaging.

### Hemolysis assays

*S. pneumoniae* strains were grown at 37°C with 5% CO_2_ until reaching mid-logarithmic phase. Bacteria were washed and a 100 µL suspension was combined with 100 µL red blood cells (isolated from buffy coat as previously described (47–49) and the mixture was incubated in a 96-well plate (Thermo Scientific^®^) at 37°C with slight agitation (300 rpm) for 30 min (positive control was only added 10 min prior to the end of the incubation). PBS was used as negative control and bi-distilled water as positive control for erythrocyte lysis. Erythrocytes derived from 9 different volunteers were tested. Then the plates were centrifuged (400g, 15min, 4°C), the supernatant was transferred to a new 96-well plate (Thermo Scientific^®^) and hemoglobin release was quantified at OD_540nm_. Statistical analysis was performed using Prism version 9 for Windows (GraphPad Software, La Jolla, CA).

### Bacterial incubation with human endothelial and epithelial cells

Endothelial HUVEC cells and epithelial A549 cells were seeded on a 18 mm diameter glass coverslips in a 12-well plate (Thermo Scientific^®^), at a concentration of 200,000 cells/well and they were grown at 37°C with 5% CO_2_ until confluence was reached. The confluent cells were then transferred to antibiotic-free growth medium and infected with pneumococci using a multiplicity of infection (MOI) of 50 bacteria per cell. The mixture was incubated at 37°C with 5% CO_2_. After washing with PBS, human cells and bacteria were fixed with 4% paraformaldehyde for 10 min at 4°C followed by blocking with 1% BSA for 1h at room temperature. A rabbit anti-*S. pneumoniae* antibody (Abcam®) was added, in order to visualize attached extracellular bacteria and also proliferating bacteria for 16h at 4°C. For HUVEC cells, concomitant incubation of a secondary anti-rabbit antibody with 4′,6-Diamidino-2-Phenylindole, dihydrochloride (DAPI, Biotium®) and Platelet endothelial cell adhesion molecule 1 (PECAM-1) conjugated with FITC was carried out for 1h at room temperature. After final washing steps with PBS, the coverslips were embedded in SlowFade Diamond (Thermo Fisher®), sealed with nail polish and stored at 4°C for subsequent imaging. Images were taken on a confocal laser scanning microscope (LSM710, Zeiss®).

### Cell viability assay

Cytotoxicity of *S. pneumoniae* D39 or the isogenic mutants towards human epithelial cells was accessed using a CellTiter-Blue® (CTB) Cell Viability Assay (Promega), according to manufacturer instructions. Human cells were seeded on a 96-well plate (Thermo Scientific^®^) with a concentration of 15,000 cells/well. Cells were cultivated at 37°C with 5% CO_2_ until confluence was reached. Then bacteria were added and the mixture was incubated for 1h under the same growth conditions. Subsequently unbound bacteria were removed by washing with DPBS. The extracellular and human cell bound pneumococci were killed by treatment of the cells with gentamicin (500 µg/mL) for 1h at 37°C under 5% CO_2_. CTB (100 μl) was added to each well. Following incubation for 16h at 37 °C in 5% CO_2_, the absorbance was measured using a Tecan® Safire 2 microplate reader at an absorption of 570 nm. In this assay, intact metabolically active endothelial cells can convert the redox dye (resazurin) into a fluorescent end product (resorufin). Statistical analysis was performed using Prism version 9 for Windows (GraphPad Software, La Jolla, CA).

### Construction of protein-protein interaction (PPI) network

The Search Tool for Retrieval of Interacting Genes (STRING) (https://string-db.org) database, which integrates both known and predicted PPIs, was applied to predict functional interactions of *S. pneumoniae* proteins (50). First, this interaction tool was used to evaluate the interaction of LytA, PspA and PspC. Second, active interaction sources, including text mining, experiments, databases, co-expression, neighborhood, gene fusion and co-occurrence and an interaction score > 0.4 were applied to construct the PPI networks. STRING is a database of known and predicted protein-protein interactions. Given a list of the proteins as input, STRING can search for their neighbor interactors and generate the PPI network consisting of all these proteins and all the interactions between them. The interactions include direct (physical) and indirect (functional) associations; they stem from computational prediction, from knowledge transfer between organisms, and from interactions aggregated from other (primary) databases (51, 52).

## Results

### LytA prevents bacterial survival within mature-biofilm structures

To explore whether and how PspA, PspC and LytA contribute to biofilm formation as the first step of adhesion, *S. pneumoniae* strains lacking the *pspA*, *pspC* or *lytA* genes were grown in polystyrene multi-well plates, and biofilm formation was quantified. LytA-lacking bacteria produced significantly more biofilm mass, whereas biofilm formation by the *pspA* and *pspC* mutants was comparable to that by the reference strain (**Fig. 1A**). The *lytA* mutant was complemented with recombinant LytA (Supplementary Fig. S1) and biofilm levels were restored to WT level, which proves the specific role of LytA in biofilm formation. To evaluate the structure of the newly formed biofilms, and bacterial survival within the tri-dimensional structure, we quantified bacterial viability using CLSM. Within the biofilms, most *lytA*-lacking bacteria were viable, as shown by green fluorescence, whereas most bacteria derived from the *pspA* and *pspC* knockout strains and WT bacteria were dead, as revealed by red fluorescence (**Fig. 1B**), which goes in accordance with the absence of lytic phase of the *lytA*-mutant. Thus, absence of *lytA* initially affects bacterial lysis and then biofilm formation is altered, whereas absence of either *pspA* or *pspC* does not influence either biofilm formation or bacterial viability.

**Figure 1.**
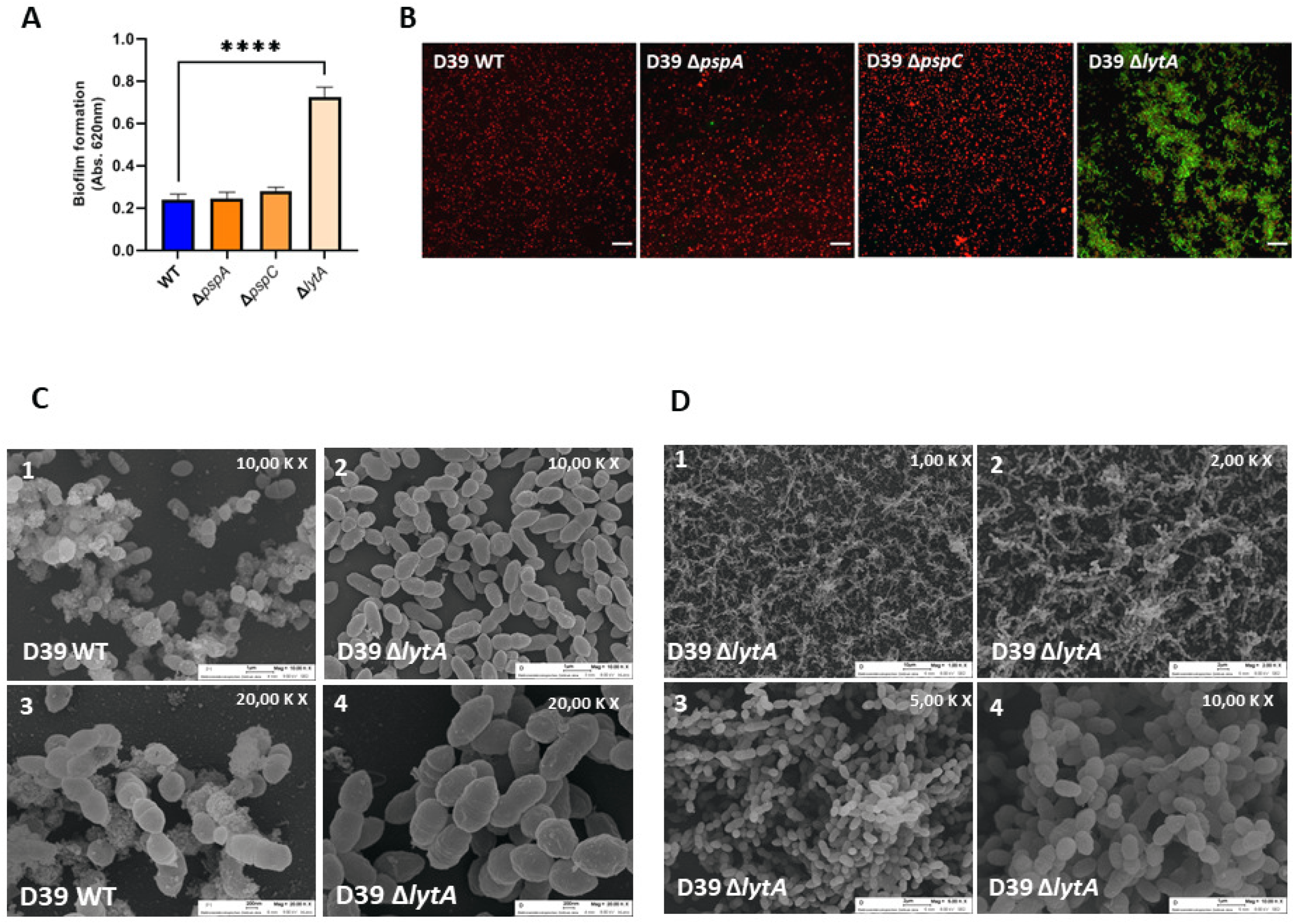
Comparison of biofilm formation among different *S. pneumoniae* strains. (A)- Quantification of biofilms grown on multi-well plates via crystal violet staining. Brown-Forsythe ANOVA test with Dunnett’s T3 multiple comparisons test and ρ< 0,0001(****). (B)- Representative images of live/dead staining on biofilms grown in glass coverslips. Red color represents dead and green alive bacteria. Scale bar is 10 µm. (C)- Representative SEM images of WT and *lytA* mutant biofilms formed on glass coverslips, at 10,000x (panel 1 and 2) or 20,000x magnifications (panels 3-4) (D)- Representative SEM images of *lytA* mutant with increasing magnifications (panel 1 to 4).

Next, we used SEM to evaluate matrix formation by the *lytA* mutant and the reference strain D39 in more detail. The *lytA* mutant produced less extracellular matrix than the parental strain D39 (**Fig. 1C**). At both low (**Fig. 1C, panel 1 and 2**) and high magnifications (**Fig. 1C, panel 3 and 4**), the WT strain (but not the *lytA* mutant) generated a prominent granular matrix. Furthermore, the *lytA* mutant generated filament-like biofilms (**Fig. 1D, panel 1 and 2**) that allowed dense bacterial agglomeration (**Fig. 1D, panel 3 and 4**). To exclude an effect of growth medium, we compared growth in THY and DMEM (**Supplementary Fig. S2**). When grown in DMEM, the Δ*lytA* strain did not produce extracellular matrix; biofilm formation was strongly impaired as visualized by lower cell numbers and the clear background; and bacterial cells were rounder, suggesting deregulated cell division. Taking into consideration the differential morphology and growth profile of Δ*lytA* strain, these results suggest an impaired extracellular matrix production on *lytA* deletion background.

### PspA, PspC and LytA reduce metabolic activity of epithelial cells

To examine the role of the three bacterial surface proteins in host cell damage, we first asked whether the mutants affect the metabolism of human alveolar epithelial cells (A549). To this end, A549 cells were co-cultivated with either knockout or WT D39 bacteria, and cell metabolism was evaluated by measuring conversion of resazurin to the fluorescent product resorufin, which only occurs in metabolically active cells. Upon contact with each of the three mutants (Δ*pspA*, Δ*pspC* and Δ*lytA*), metabolic activity in cells increased, and was higher than in cells challenged with the wildtype strain D39 (**Fig. 2A**). This shows that the pathogenic reference strain can damage human epithelial cells and therefore decrease metabolic activity of human epithelia is observed, and that deletion of a single CBP gene affects the ability to cause cell damage. Each mutant showed different effects. The Δ*lytA* strain affected cell metabolism more strongly than Δ*pspA* or Δ*pspC*. Thus, deletion of one single CBP affects the metabolic turnover of human epithelial cells, thereby confirming that each CBP contributes to pathogenicity.

**Figure 2.**
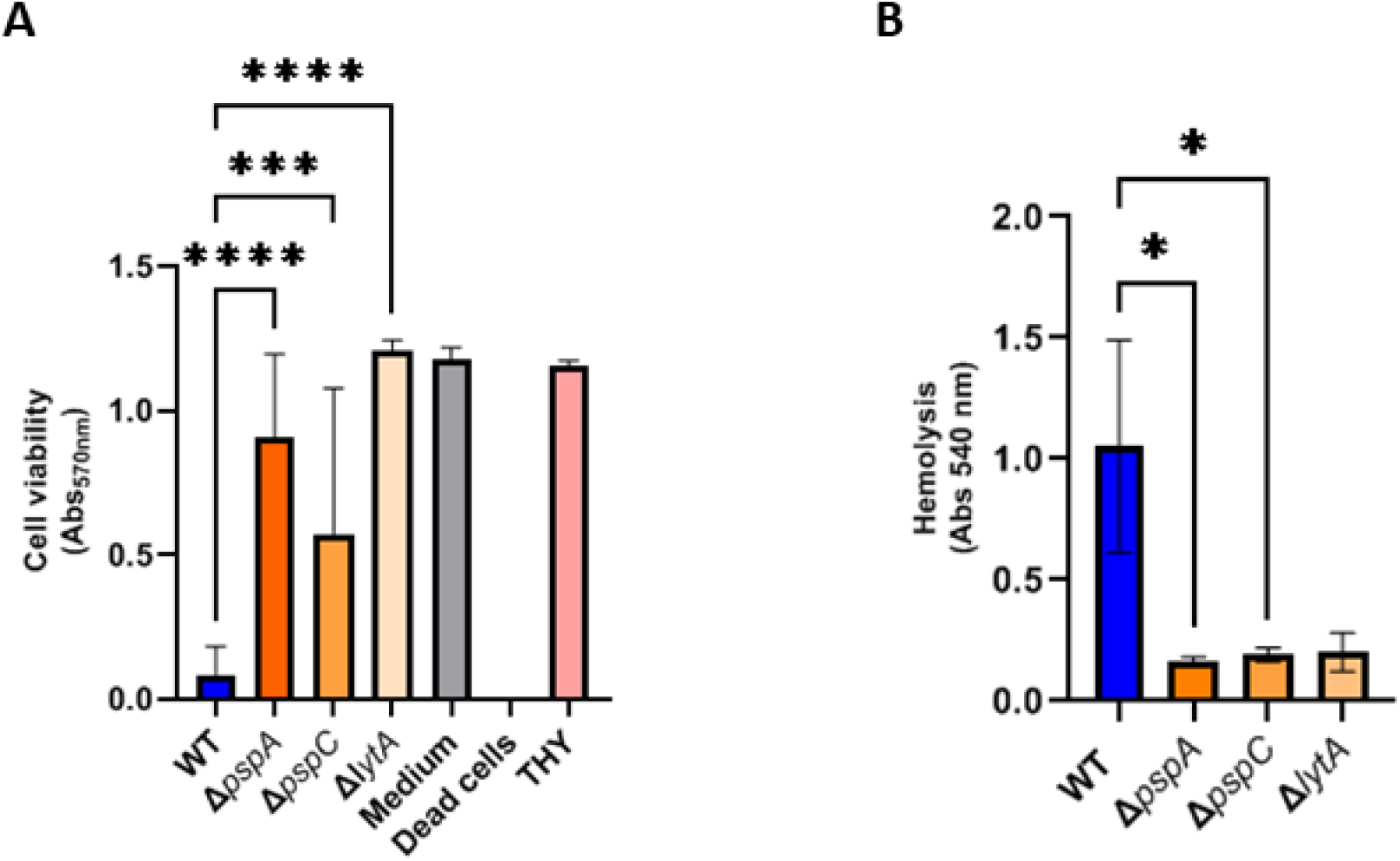
Evaluation of the effect of different *S pneumoniae* strains on red blood and epithelial cells. (A)- Assessment of A549 cell viability by cell cytotoxicity assay. Human epithelial cells were challenged with either WT (*S. pneumoniae* D39) or the indicated mutant strains. DMEM, THY and dead A549 cells were used as controls. One-way ANOVA with Dunnett’s multiple comparisons test and ρ<0,0001(****),ρ<0,0005(***). (B)- Hemolysis of human red blood cells by *S. pneumoniae* strains. One –way ANOVA with Tukey’s multiple comparisons test and ρ<0,05(*).

### The three CBPs mediate pneumococcal hemolytic activity

To further define the role of the three CBPs on interactions with host cells, we evaluated the haemolytic capacity of the knockout mutants. The *S. pneumoniae* mutants were added to human red blood cells and, following incubation, erythrocyte lysis was evaluated. Erythrocyte lysis induced by each knockout was less than that induced by the pathogenic reference strain D39 (**Fig. 2B**). Thus, each CBP contributes to the ability of the bacteria to induce erythrocyte lysis.

### The three CBPs induce expression of PECAM-1 by endothelial cells

After addressing biofilm formation and the effects on red blood cells, we next examined the effects of the bacterial mutants on human endothelial cells (HUVECs). Expression of the human endothelial cell surface marker platelet-endothelial cell adhesion molecule-1 (PECAM-1) (53–55) was monitored by CLSM. HUVEC cells were cultivated with either the CBP mutants or the WT strain. Each knockout mutant strongly reduced surface expression of PECAM-1 (**Fig. 3A**). Moreover, Δ*lytA* strain sustained bacterial growth when in contact with human endothelial cells, as seen by the intense red fluorescence signal (bacteria) in the Δ*lytA* panel; however, it did not induce expression of this surface marker. Semi-quantitative analysis of the microscopy data showed significant upregulation of PECAM-1 expression only in the presence of the reference D39 strain (**Fig. 3B**), corroborating the observed absence of cellular damage upon incubation with any of the CBP mutants. The pathogenic strain D39 was efficiently internalized by HUVEC cells, but all mutant strains showed less internalization by the human cells (**Fig. S3**). Thus corroborating a lower adhesion and recognition of the mutants strains, lower levels of PECAM-1surface expression and less internalization by endothelial cells.

**Figure 3.**
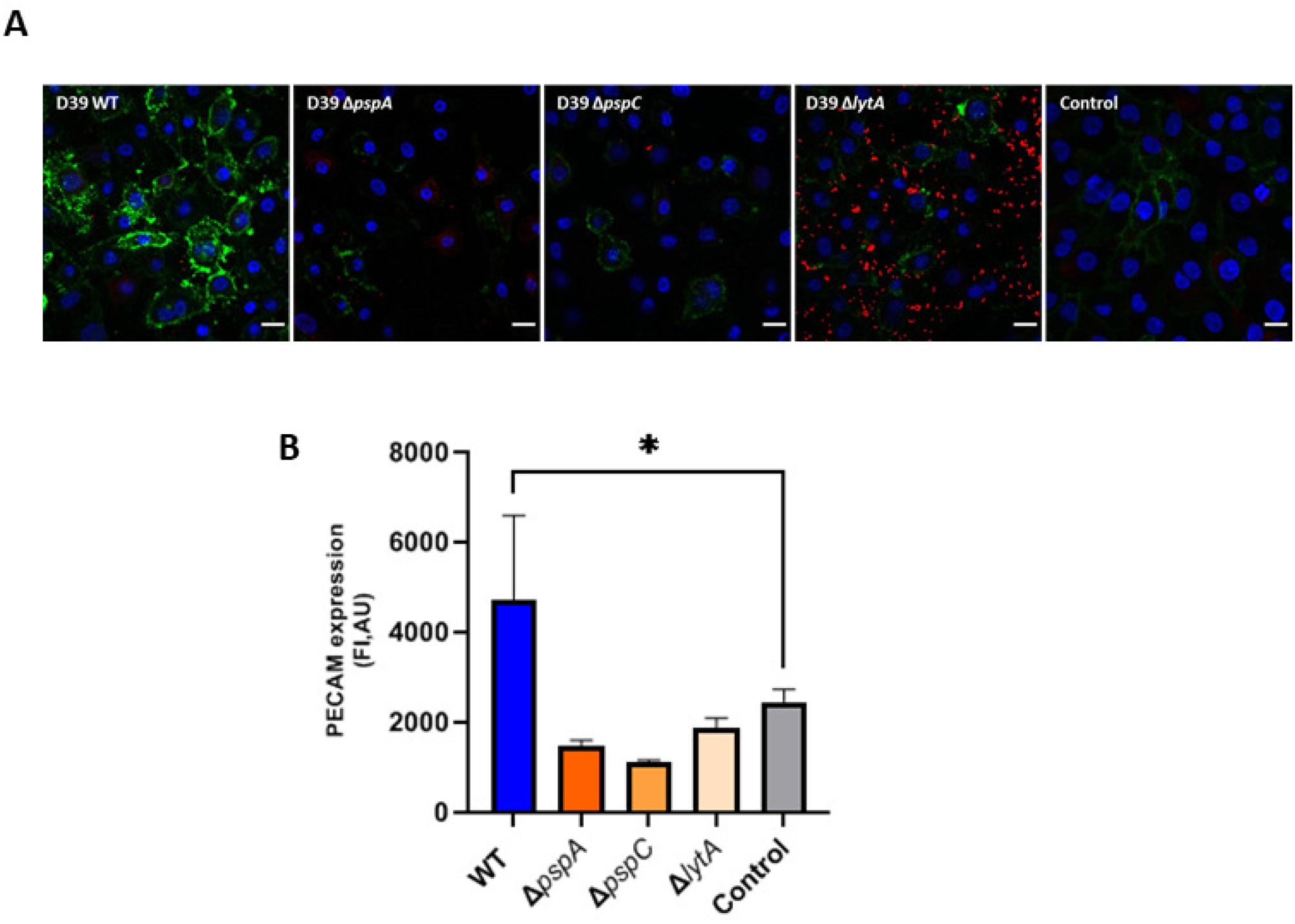
Evaluation of the effect of different *S pneumoniae* strains on human endothelial cell surface marker expression. (A)- Representative images of HUVEC-bacteria co-incubation visualized by CLSM. PECAM-1 FITC signal is represented in green, nuclei are stained in DAPI (blue) and bacteria are stained with anti-pneumococcus antiserum (red fluorescence). Scale bar 10µm. (B)- Quantification of PECAM-1 expression. One-way ANOVA with Dunnett’s multiple comparisons test and ρ<0,05(*).

**Table 1** summarizes the effects of each pneumococcal mutant and the reference D39 strain. Apart from the effect of Δ*lytA* on biofilm formation, all other effects suggest that mutants show impaired pathogenic capacity, suggesting that the corresponding proteins play a crucial role in evading host immune responses.

**Table 1.** Summarizing table of the influence of each strain on pneumococcal pathogenic features. Upwards arrow indicates an increase activity, downwards arrow indicate a decrease activity.

| **Feature** | **D39** | ***ΔpspA*** | ***ΔpspC*** | ***ΔlytA*** |
| --- | --- | --- | --- | --- |
| Biofilms | ↓ | ↓ | ↓ | ↑ |
| Epithelial metabolism | ↓ | ↑ | ↑ | ↑ |
| Haemolysis | ↑ | ↓ | ↓ | ↓ |
| Endothelial PECAM-1 expression | ↑ | ↓ | ↓ | ↓ |

| Feature | D39 | $\Delta pspA$ | $\Delta pspC$ | $\Delta lytA$ |
| --- | --- | --- | --- | --- |
| Biofilms | ↓ | ↓ | ↓ | ↑ |
| Epithelial metabolism | ↓ | ↑ | ↑ | ↑ |
| Haemolysis | ↑ | ↓ | ↓ | ↓ |
| Endothelial PECAM-1 expression | ↑ | ↓ | ↓ | ↓ |

### PspA, PspC and LytA are part of a complex network that regulates host–pathogen interactions

So far, the results show that each pneumococcal protein plays multiple roles during host–pathogen interactions, i.e., biofilm formation, haemolysis, and epithelial and endothelial cell damage. These effects are distinct and involve diverse pneumococcal cellular subsystems, such as the cell division machinery (to regulate growth rate), expression of toxins (to induce host cell damage), and peptidoglycan synthesis (to release haemolytic enzymes). Therefore, we evaluated the connections between the three proteins within protein networks.

We constructed protein-protein interaction (PPI) networks for each pneumococcal protein using the STRING database. The initial network focused on PspA, PspC and LytA, and showed that LytA mediates the interaction between PspA and PspC (**Fig. 4A**). An enlarged and more complex network map was then constructed based on several criteria; strong connections are represented by thick, dark-grey lines whereas weaker connections (with fewer matching criteria) are represented by shaded, light-grey lines (**Fig. 4B**). The annotated proteins are presented next to the circular intersection node icon. After PPI construction, a Markov Cluster Algorithm (MCL) was used to identify cluster-specific groups (56). Four protein clusters appeared: a very densely interconnected cluster I (red), a small yet strongly interconnected cluster II (brown), cluster III comprising connections with different degrees of strength (green), and cluster IV (yellow) representing fewer interacting partners. Each cluster contained (predominantly) members related to a specific cellular machinery: cluster I = cell division; cluster II = chromosome replication; cluster III = peptidoglycan biosynthesis; and cluster IV = unannotated proteins.

**Figure 4.**
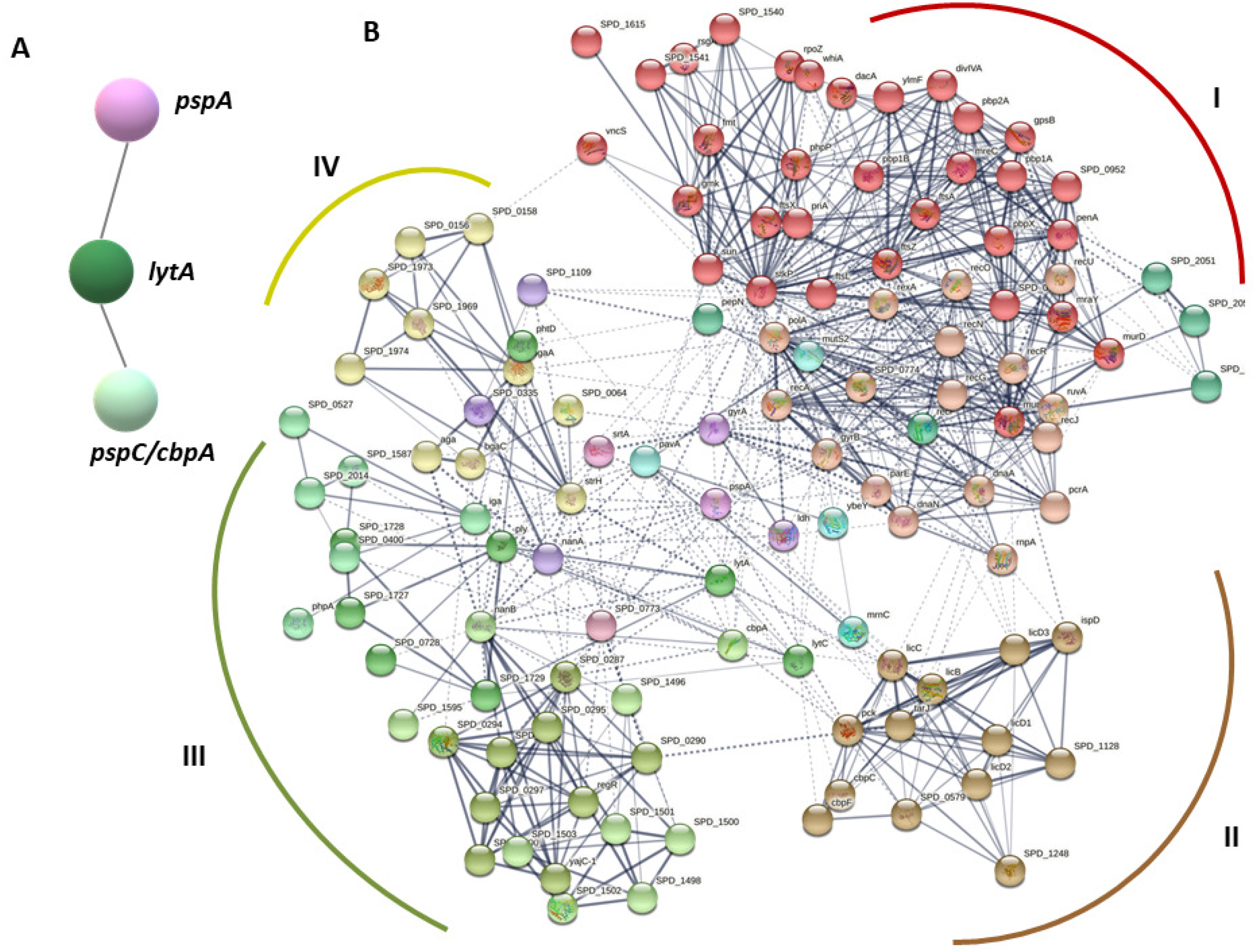
STRING Protein-Protein Interaction (PPI) Networks. (A)- Linear interaction network of the three studied CBPs, LytA, PspA and PspC (CbpA). Colorful circles or nodes represent the input proteins. (B)- PPI of interaction partners of PspA, PspC (CbpA) and LytA. Strength (confidence) of interaction is represented by thicker grey lines. Dashed lines represent interactions based in fewer criterions. The different colors are specific of different clusters (MCL clustering).

Each protein (PspA, PspC or LytA) is embedded in a complex network of cross-interactions, and each is integrated at a very prominent position, suggesting that targeting the proteins therapeutically can destabilize intracellular bacterial homeostasis. Additionally, this prominent positioning contextualizes and connects all of the data, suggesting that these CBPs have a multifactorial effect on host invasion and cell damage.

## Discussion

*S. pneumoniae* expresses a family of CBPs that use related cell wall anchors and act as central virulence proteins that interact with the soluble host complement regulators and with human immune system. Here, we evaluate and compare three important CBPs, i.e., PspA, PspC and LytA, with respect to their effects on bacterial biofilm formation and host cell damage, and their integration into bacterial protein networks.

Pneumococci lacking LytA form more biofilm extracellular matrix. By contrast, PspA and PspC have no such effects on biofilm matrix formation. Previous studies show that CBPs play roles in invasion of the host under various environmental conditions (40, 57). Efforts have been made to evaluate the role of extracellular DNA-CBPs complexes during establishment of biofilms; interestingly, LytA-mediated DNA complexes are relevant to biofilm formation (40). Here, we show that PspA and PspC mediate biofilm formation, which supports current knowledge; however, our observations regarding LytA are rather different (**Fig. 1**). We found that LytA prevents, rather than promotes, biofilm formation, as bacteria lacking the gene encoding autolysin were more likely to form biofilms. Similar to the results presented here, another study reported an anti-biofilm effect of LytA (58); however, few studies have examined the specific and peculiar role of LytA in pneumococcal biofilms. Protein network analysis revealed that LytA is linked and integrated into the bacterial cell division machinery and teichoic acid biosynthesis (**Fig. 4**). Triggering of the bacterial SOS response cascade, which is required for establishment of biofilms, might affect transcription of the *lytA* operon, most likely leading to repression.

This study also identifies a novel role for PspA, PspC and LytA in haemolysis. Bacteria lacking *pspA*, *pspC* or *lytA* are less efficient at inducing lysis of human erythrocytes (**Fig. 2B**). The PPI network map (**Fig. 4B**) shows that pneumolysin (ply), a lytic pore forming pneumococcal enzyme, interacts closely with LytA (and even LytC, another pneumococcal CBP). However, PspA and PspC do not interact with ply. Instead, these two proteins interact with neuraminidase (NanA), a *S. pneumoniae* enzyme that is upregulated upon contact with the host cells; secreted NanA cleaves host glycoconjugates (60). This relationship between PspA, PspC and NanA might connect haemolysis and the two surface-bound pneumococcal proteins, and provide another link to biofilm formation. A recent study proposes that pneumococcus haemolysis activity is mediated mainly by hydrogen peroxide (H_2_O_2_) rather than by excreted pneumolysin (48,61,62). In terms of the haemolytic profile, the pneumococcal mutants presented in that study, which do not produce H_2_O_2_, show a surprising resemblance to the *pspA*, *pspC* and *lytA* mutants in the present study. Moreover, H_2_O_2_ produced by *S. pneumoniae* damages human lung cells (63). Therefore, the reduced cytotoxicity of the CBPs mutants toward human epithelial and endothelial cells (**Fig. 2A and 3**) might be a consequence of decreased H_2_O_2_ production.

To obtain a deeper understanding of the role of CBPs in epithelial and endothelial damage, a thorough metabolome analysis should be performed to examine the intracellular metabolic status of the mutants.

The association between bacterial cell wall- and/or membrane-anchored proteins and the stress response has been described before, with extensive studies being carried out in Gram-negative bacteria, particularly *Escherichia coli* (64–66). In Gram-positive bacteria, the thick peptidoglycan layer is responsible for protection against environmental cues. Yet, other studies show the dynamic nature and diverse structure of the peptidoglycan layer across the cell wall of even a single bacterium (67, 68). This diversity may lead to distinct distributions of bacterial surface-exposed proteins within a certain period of time, which might dictate the interaction with the human host. The interchange of information between the different intracellular metabolic pathways, and the arrangement of the cell wall (mainly the CBPs) is a crucial immune evasion strategy.

A thread connecting all of our experimental findings is the intermingled network of interactions sustained by the three CBPs (**Fig. 4B**). The PPI network map shows four clusters that relate to different cellular pathways. Cluster I contains mainly proteins related to the cell division machinery (DivIVA, FtsZ, FtsL, FtsX and GpsB) (69); cluster II contains proteins related to chromosome replication (DnaA, DnaN, ParE and PriA) (70); and cluster III contains proteins related to peptidoglycan biosynthesis (Pbp1B, Pbp2A, PenA, MurC and MurD) (71). A relevant interaction partner for each evaluated protein is the serine/threonine-protein kinase StkP (62,72–74). StkP plays a crucial role in regulating cell shape and division of *S. pneumoniae* through control of DivIVA activity. StkP is thought to sense intracellular peptidoglycan subunits in the division septum of actively growing cells and to adjust the regulation of DivIVA. The bond between StkP and the triad PspA-PspC-LytA suggests a link between each CBP in the cell wall of pneumococci and intracellular regulation subjacent to the synthesis of the cell wall itself, as well as coordination of cell division. Hence, *S. pneumoniae* fine-tunes expression of surface-anchored immune evasion proteins in response to the cell cycle. Cluster II comprises additional CBPs such as CbpC and CbpF (75), and proteins related to teichoic acid biosynthesis (LicB, LicC and LicD) (76). The large cluster III also contains regulators with different backgrounds and functions, e.g., Ply (pneumolysin) (59), NanA and NanB (both of which are implicated in biofilm formation and in colonization of the upper airway) and PavA (which is critical for the overall virulence of pneumococci) (77). Cluster IV harbours transcriptional regulators, enzymes and other uncharacterized proteins.

Understanding the interconnectivity of PspA, PspC and LytA with other subcellular systems and their impact on bacterial homeostasis, is pivotal to mechanistically comprehend immune evasion by these proteins and to developing effective therapeutics.

## Acknowledgements

This work was supported by the Collaborative Research Center, FungiNet (project C6), Deutsche Forschungsgemeinschaft (DFG). SH received funding from Deutsche Forschungsgemeinschaft (DFG HA 3125/5-2). PFZ received support from the KIDNEEDS Foundation Iowa City USA:

## Author Contributions

CV conceived and designed the experiments. CV, SD, MB and TPK performed the experiments. CV, SD and MB analysed the data. MW, SH and PZ contributed with reagents, materials, and analysis tools. CV and PZ wrote the manuscript and TPK and SH edited the manuscript.

